# Mechanistic Insights into Cyclic Penta-Adenylate-Mediated Activation of Type III CRISPR Ribonuclease Csm6

**DOI:** 10.1101/2025.10.25.684504

**Authors:** Ruyi Shi, Mengquan Yang, Yusong Liu, Haishan Gao, Zhonghui Lin

**Author notes:** Correspondence (H. G.); (Z. L.). These authors contributed equally to this work (Ruyi Shi, Mengquan Yang).

## Abstract

Type III CRISPR systems generate cyclic oligoadenylate (cOA, 3 to 6 AMPs) messengers upon detecting viral RNA, activating downstream effectors to defend against viral infection. Although cOA-activated effectors have been extensively characterized, the cA_5_-specific effectors remained unexplored despite cA_5_ being among the most abundant cOA species produced during phage infection. Here, we report that *Actinomyces procaprae* Csm6 (ApCsm6) selectively employs cA_5_ as its activator. Unlike other characterized Csm6 proteins, ApCsm6 self-limits its ribonuclease activity by degrading cOAs via its HEPN domain, rather than relaying on the CARF domain. Cryo-EM structures of ApCsm6 and its complexes with cA_5_ and cA_6_ reveal a homotetrameric assembly, where each monomer binds a single cOA within a composite pocket formed by two tandem CARF-HEPN domains. Binding of cA_5_, but not cA_6_, enhances tetramerization and induces large conformational shifts in CARF, which in turn allosterically activates ssRNA cleavage in HEPN. These findings advance our understanding of ligand discrimination and signaling regulation in type III CRISPR immunity.

**Highlights:**

- ApCsm6 preferentially recognizes cA_5_ as its activator.
- Cryo-EM structures of ApCsm6 and its complexes with cA_5_ or cA_6_ reveal the structural basis for cA_5_-selective recognition and allosteric activation.
- ApCsm6 acts as a self-limiting ribonuclease by degrading cOAs via its HEPN domain rather than relaying on the CARF domain.

## Introduction

Bacteria and archaea employ CRISPR-Cas systems to defend against invading foreign genetic elements ^1–3^. Unlike other CRISPR-Cas systems, type III systems are distinguished by a unique cyclic oligoadenylate (cOA) signaling mechanism, which provides an additional defense against viral RNA ^4–11^. Upon detecting viral mRNA, the Cas10 protein (also referred to as Csm1 or Cmr2) within type III CRISPR-Cas complexes synthesizes ring-shaped cOA molecules, which are composed of three to six AMPs linked by 3’-5’ phosphodiester bonds ^4–10^. These messenger cOAs in turn can activate diverse downstream effector proteins, including RNases, DNases, transcription factors, proteases as well as adenosine deaminase ^10,12–16^.

To date, the most characterized type III CRISPR effectors are the RNA-targeting Csx1/Csm6 proteins. These proteins typically consist of an N-terminal CARF (CRISPR-associated Rossman-fold) domain and a C-terminal HEPN (higher eukaryotes and prokaryotes nucleotide) domain ^16–21^. When cOA binds to the CARF domain, it allosterically activates ribonuclease activity in the HEPN domain, resulting in nonspecific degradation of ssRNA ^20–22^. To prevent excessive RNA degradation that could cause host cell dormancy or death during early viral infection, cells utilize a group of cOA-degrading enzymes called ring nucleases ^23–30^. Beyond the standalone ring nucleases, certain Csx1/Csm6 proteins, referred to as self-limiting ribonucleases, exhibit intrinsic cOA-degrading activity. This activity is mediated either through their own CARF domains ^19,20,22,24,31,32^ or via the integration of a viral anti-CRISPR ring nuclease ^21,33^, providing an off-switch to downregulate cOA signaling.

Although the Cas10 nucleotidyl cyclase generates a spectrum of cOAs (including cA_3_, cA_4_, cA_5_, and cA_6_), most characterized type III effectors prefer cA_4_ as their cognate ligand, with a subset utilizing cA_6_ ^10,12,13^. This preference is likely attributed to the homodimeric assembly of their CARF domains, which favor recognition of ligands with twofold symmetry. Likewise, the CBASS (cyclic oligonucleotide-based anti-phage signaling system) effector NucC recognizes cA_3_ as a homotrimer ^34^. In contrast, the distant CARF homologs, SAVED (SMODS-associated and fused to various effector domains) domains function as monomers and bind a range of cyclic di- and trinucleotides ^10^. For example, the SAVED domain of Cap4 comprises two tandem CARF-like subdomains and specifically recognizes cA_3_ as its activator ^35^. Previous bioinformatic analysis identified a Csm6 homolog from *Actinomyces procaprae*, designated Csm6-2, which consists of two tandem CARF-HEPN modules in a single polypeptide chain ^36^. Csm6-2 presents in 16 type III-D loci and is hypothesized to have originated from a Csm6 ancestor via gene fusion ^36^, thereby representing an uncharacterized class of cOA-responsive CARF effectors.

In the present study, we show that ApCsm6 preferentially employs cA_5_ as its activator. In contrast to other characterized Csm6 proteins, ApCsm6 uniquely employs its HEPN domain (rather than the CARF domain) to degrade cOAs, enabling self-limitation of its ribonuclease activity. Furthermore, the cryo-EM structures of ApCsm6 and its complexes with cA_5_ and cA_6_ reveal the molecular basis for cA_5_-specific recognition and allosteric activation of ssRNA cleavage. These findings advance our understanding of type III CRISPR signaling systems, providing new insights into ligand specificity and regulatory mechanisms.

## Results

### ApCsm6 selectively recognizes cA_5_ as its Activator

We first characterized ApCsm6’s ribonuclease activity in response to various cOAs (cA_4_, cA_5_, and cA_6_). In a gel-based assay, while cA_4_ showed no activation of ssRNA cleavage, cA_6_ stimulated detectable activity (Fig. 1a). Notably, under identical conditions, cA_5_ induced robust ribonuclease activity (Fig. 1a). Quantitative analysis using a FRET-based cleavage assay revealed that cA_5_ activates ApCsm6 with an EC_50_ of 0.09 nM, demonstrating ∼1,000-fold greater potency than cA_6_ (EC_50_ = 103.3 nM) (Fig. 1b).

**Fig. 1.**
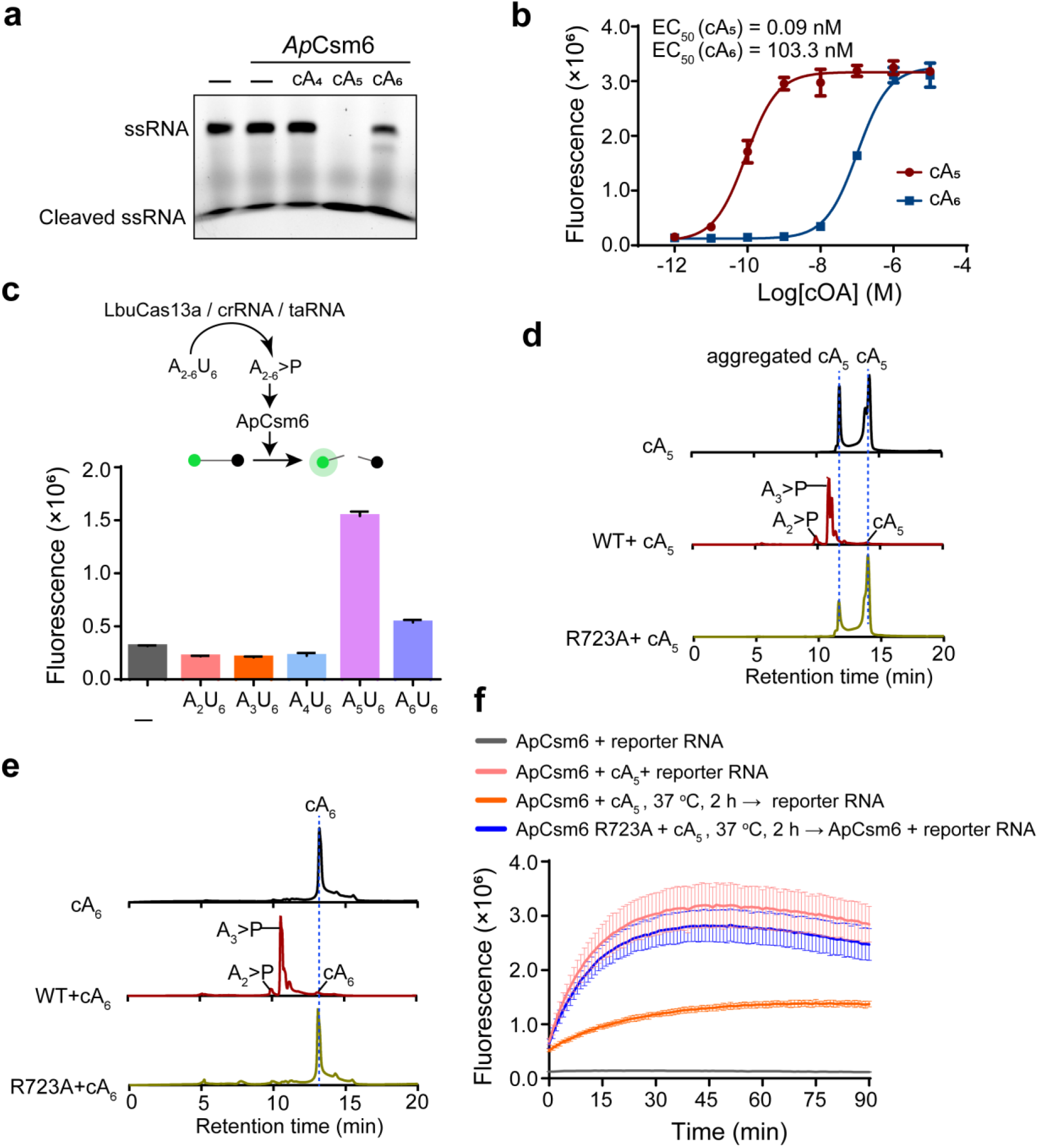
cA5 is the preferred activator of ApCsm6. a, Effect of various cOAs on the stimulation of ssRNA cleavage by ApCsm6. 100 nM ApCsm6 protein was incubated with 100 nM cOAs and 250 nM FAM-labeled ssRNA at 37°C for 45 min, followed by denatured polyacrylamide gel analysis. **b,** Dose-responsive curves for the activation of ApCsm6 by cA_5_ and cA_6_ in the FRET-based ssRNA cleavage assay. Values are means ± SD, n = 3 replicates. **c,** LbuCas13a-Csm6 tandem assay assessing linear oligonucleotide activation. Top: Assay schematic. Bottom: FRET-based cleavage with 40 nM LbuCas13a, 20 nM crRNA, 100 pM target RNA, 100 nM ApCsm6, 200 nM reporter RNA, and 2 μM A_2-6_U_6_ activator. Values are means ± SD, n = 3 replicates. **d,e,** HPLC analysis of cOA degradation by WT and mutant ApCsm6. Reactions were conducted at 37°C for 2 h, using 40 μM cOA and 2 μM ApCsm6. **f,** FRET-based ssRNA cleavage assay showing reduced ribonuclease activity after pre-incubation of 1 nM ApCsm6 with 10 nM cA_5_. Values are means ± SD, n = 3 replicates.

To further characterize cOA activation profiles, we employed a LbuCas13a-Csm6 tandem nuclease assay that leverages LbuCas13a’s collateral ribonuclease activity upon target RNA detection ^37^ (Fig. 1c). This activity processes the poly (U) region of A_2-6_U_6_ to generate linear oligoadenylate activators containing 2′,3′-cyclic phosphates (A_2-6_>P) that exhibit activation potency comparable to their cyclic counterparts ^37^. Consistent with the above results, A_5_>P demonstrated significantly stronger activation of ApCsm6-mediated ssRNA cleavage compared to A_6_>P (Fig. 1c). In contrast, A_2_>P, A_3_>P, and A_4_>P failed to elicit any detectable activation (Fig. 1c).

Together, these findings establish cA_5_ as the preferential activator of ApCsm6, distinguishing it from other characterized Csm6 proteins that rely on cA_4_ or cA_6_.

### ApCsm6 functions as a self-limiting ribonuclease by degrading cOA activators through its HEPN domain

As most Csm6 proteins function as self-limiting ribonucleases through intrinsic cOA degradation, we sought to determine whether ApCsm6 shares this capability. HPLC and MALDI-TOF MS analyses revealed that incubation with 2 μM ApCsm6 at 37°C for 2 h degraded both cA_5_ and cA_6_, generating predominantly A_2_>P and A_3_>P products (Fig. 1d, e and Supplementary Fig. 1). Notably, the R723A mutation in the HEPN domain completely abolished this activity, indicating that ApCsm6 degrades cOAs through its HEPN domain (Fig. 1d, e).

To determine whether cOA degradation regulates ApCsm6’s ssRNA cleavage activity, we pre-incubated cA_5_ with ApCsm6 at 37°C for 2 h before adding ssRNA. This treatment reduced ssRNA cleavage by approximately 70% compared to non-pre-incubated control (Fig. 1f), demonstrating that cOA turnover negatively regulates ApCsm6 function. Importantly, the R723A mutation nearly eliminated this regulation (Fig. 1f), confirming the role of HEPN domain in the self-limiting regulation.

Collectively, these results suggest that ApCsm6 functions as a self-limiting ribonuclease through HEPN domain-mediated degradation of its cOA activators.

### Overall architecture of ApCsm6

The full-length ApCsm6 protein comprises two tandem CARF-HEPN domains: CARF1-HEPN1 (residues 1–400) and CARF2-HEPN2 (residues 401–797) (Fig. 2a). To elucidate how the two CARF-HEPN modules assemble into a cOA-responsive ribonuclease, we determined the cryo-EM structure of ApCsm6 in its apo state. The structure was resolved at a resolution of 2.59 Å (Supplementary Fig. 2 and Supplementary Table 1). It reveals that ApCsm6 assembles as a homotetramer (Fig. 2b, c). Within each monomer, the CARF1-HEPN1 and CARF2-HEPN2 modules are structurally integrated, forming two composite domains, CARF and HEPN, for cOA-binding and ssRNA cleavage, respectively (Fig. 2d). Unlike other homodimeric Csm6 proteins, which adopt a symmetric “X”-shape, ApCsm6 shows a pronounced tilt of its CARF domain relative to the HEPN domain (Fig. 2d).

**Fig. 2.**
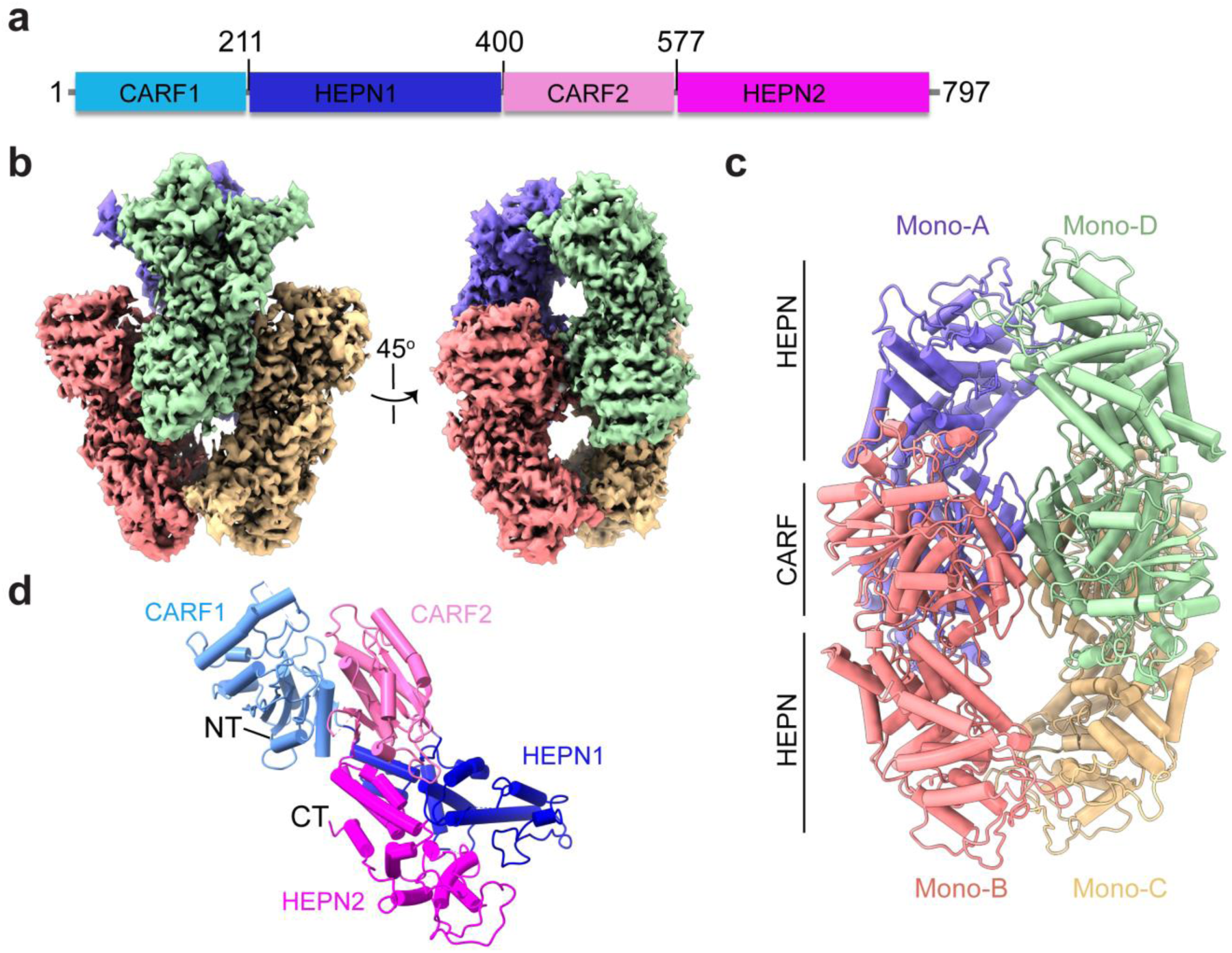
Cryo-EM structure of ApCsm6. a, Schematic diagram of the domain architecture of ApCsm6. **b,** Cryo-EM map for ApCsm6, with each monomer shown in a distinct color. **c,** Cartoon representation of ApCsm6. **d,** Structure representation of an ApCsm6 monomer. Subdomains are colored as in (a).

The tetramerization of ApCsm6 is primarily mediated by CARF2 domain (Fig. 3a, b). First, two pairs of ApCsm6 monomers (A/B and C/D) assemble into dimers via a pair of antiparallel β-strands and helices (Fig. 3c). These dimer interfaces are stabilized by a network of hydrogen bonds and hydrophobic interactions involving residues V466, L473 and I488 (Fig. 3c). Then, two such dimers further associate into a tetramer mainly through four adjacent α-helices, mainly mediated through hydrogen bonds between K467 and T476, as well as hydrophobic interactions involving L473 and L474 (Fig. 3d, e). Mutation of L473 and L474 to glutamines disrupted tetramer formation (Fig. 3f, g), leading to a significant reduction in ssRNA cleavage activity (Fig. 3h). Collectively, these results indicate that ApCsm6 functions as a homotetramer.

**Fig. 3.**
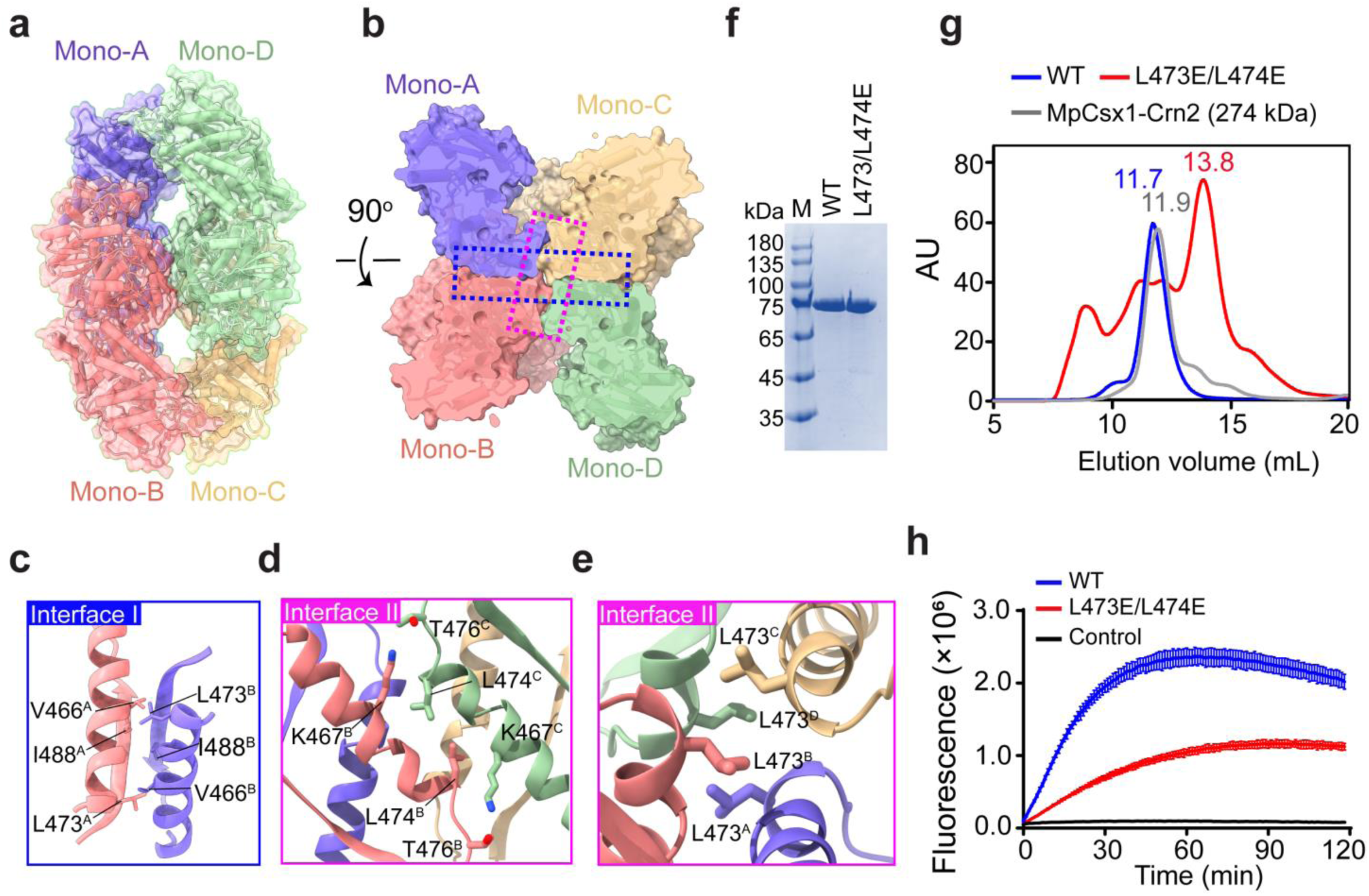
ApCsm6 functions as a homotetramer. a,b, Structure of ApCsm6 overlaid with 60-80% transparent surface. **c-e,** Close-up views of the interfaces between monomers. **f,** Coomassie blue-stained gel of recombinant wild-type (WT) and mutant ApCsm6 proteins. **g,** Size-exclusion chromatography profiles of the indicated proteins. **h,** Ribonuclease activity of WT and mutant ApCsm6, measured using a FRET-based ssRNA cleavage assay. 200 nM ApCsm6 proteins were incubated with 200 nM cA_6_ and 200 nM FAM/BHQ1-labeled ssRNA at 37°C. Fluorescence intensities were recorded at 1-min intervals. Values are means ± SD, n = 3 replicates.

### Structure of ApCsm6 in complex with cA_6_

To investigate the mechanism of cOA-dependent activation, we next sought to determine the structure of ApCsm6 in complex with cOA. To prevent cOA cleavage by the HEPN domain of ApCsm6, we introduced a R723A mutation. The structure of ApCsm6-cA_6_ complex was solved at a resolution of 2.53 Å (Supplementary Fig. 2 and Supplementary Table 1). It reveals a homotetrameric assembly of ApCsm6 as observed in its apo form. Each monomer binds a single cA_6_ molecule within a composite pocket formed by two tandem CARF-HEPN domains (Fig. 4a). The cA_6_ molecules exhibit well-resolved electron density, except for one AMP (AMP-6), which has poorly-defined density (Fig. 4b). Within the cA_6_ ring, four AMP moieties (AMP-1, 2, 5 and 6) are bound within CARF1, while the remaining two (AMP-3 and -4) are localized to CARF2 (Fig. 4c, d).

**Fig. 4.**
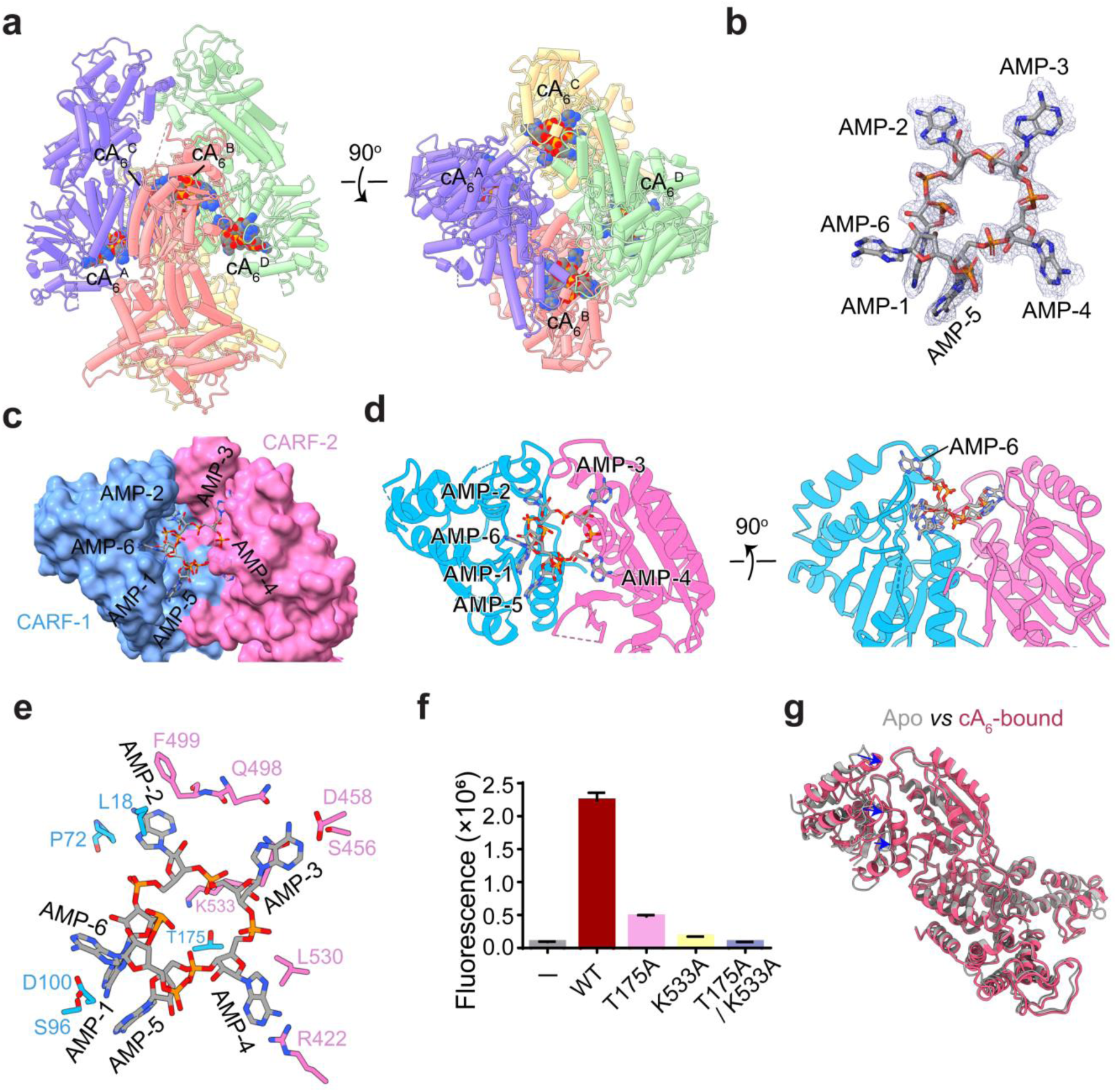
Structure of ApCsm6 in complex with cA_6_. a, Cartoon representation of ApCsm6-cA_6_ complex. α-helices are represented as cylinders, and cA_6_ is in sphere. **b,** Stick representation of cA_6_ overlaid with its cryo-EM map. **c,d,** Surface and cartoon diagrams of the ApCsm6 CARF domain bound to cA_6._ The CARF-1 and CARF-2 moieties are colored in blue and pink, respectively. **e,** Close-up view of interactions between cA_6_ and the ApCsm6 CARF domain. **f,** Effect of CARF domain mutations on the ribonuclease activity of ApCsm6. Values are means ± SD, n = 3 replicates. **g,** Structural comparison of the apo (grey) and cA_6_-bound (red) ApCsm6. Structural changes upon cA_6_ binding are indicated by blue arrows.

Similar to other cOA-dependent type III CRISPR effectors, the phosphodiester ring of cA_6_ localizes onto the top of the interface between CARF-1 and CARF-2 interface (Fig. 4c,d). In particular, residues K533 and T175 project their side chains into the ring center, coordinating the 5’-phosphates of AMP-2 and AMP-5, respectively. Alanine substitution of T175 or K533 significantly reduced ApCsm6’s ribonuclease activity, while mutation of both residues abolished activity (Fig. 4f). The adenine groups of AMP-1 to AMP-4 are each accommodated within a specific pocket via extensive hydrogen bonds and hydrophobic interactions (Fig. 4e). By contrast, AMP-5 is stabilized primarily through base-stacking interactions with AMP-1, while AMP-6 makes minimal contact with the protein (Fig. 4e).

The overall structure of cA_6_-bound ApCsm6 closely resembles that of its apo form, with minor conformational changes in the CARF-1 domain, which shifts slightly towards the CARF2 domain upon cA_6_ binding (Fig. 4g and Supplementary Movie 1).

### Mechanism of specific cA_5_ recognition

To illuminate the mechanism of ApCsm6’s preference for cA_5_ as its activator, we determined a 2.67 Å cryo-EM structure of cA_5_-bound ApCsm6 harboring a HEPN-inactivating mutation H369A (Supplementary Fig. 2 and Supplementary Table 1). The structure reveals that ApCsm6 adopts an overall architecture similar to its cA_6_-bound form, with each monomer binding a single cA_5_ molecule (Supplementary Fig. 3). Strikingly, unlike the apo or cA_6_-bound states, the CARF domain of cA_5_-bound structure undergoes a large conformational change, transitioning to a closed state that nearly encapsulates the entire cA_5_ ligand (Fig. 5a). All five AMP moieties of each cA_5_ molecule exhibit well-defined electron density (Fig. 5b), but cannot be fully aligned with those of cA_6_ (Fig. 5c).

**Fig. 5.**
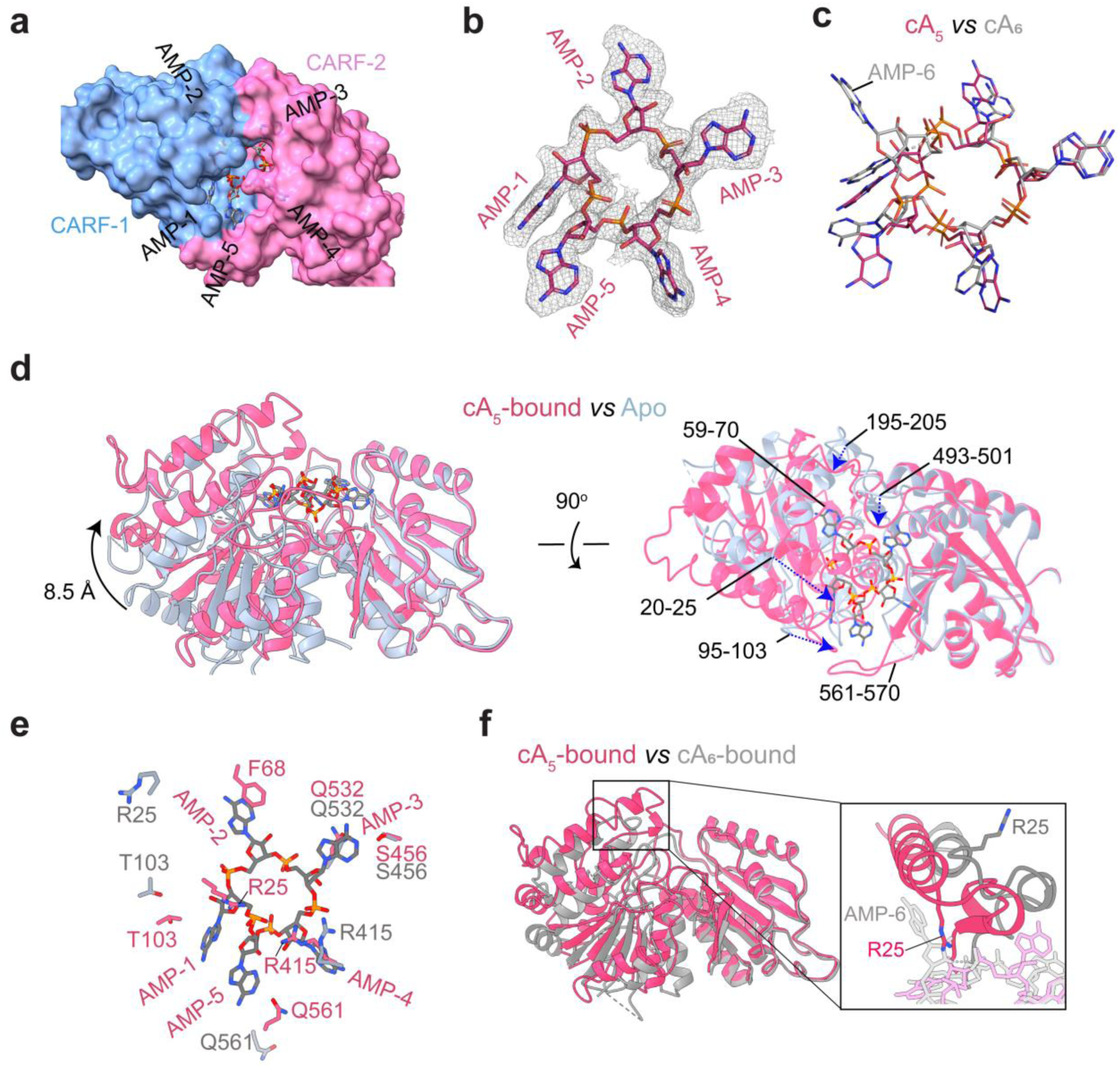
Structure of ApCsm6 in complex with cA_5_. a, Surface representation of the ApCsm6 CARF domain bound to cA_5._ **b,** Stick representation of cA_5_ overlaid with its cryo-EM map. **c,** Superimposition of cA_5_ (magenta) and cA_6_ (grey). **d,** Structural alignment of the cA_5_-bound ApCsm6 CARF domain (pink) with its apo form (light blue). Major conformational changes upon cA_5_ binding are labeled and indicated by blue arrows. **e,** Close-up view of interactions between cA_5_ and the ApCsm6 CARF domain. Residues involved in cA_5_ recognition are shown as sticks, and compared with their corresponding residues in the apo state (light blue). **f,** Structural alignment of the cA_5_-bound ApCsm6 CARF domain (pink) with the cA_6_-bound form. The zoomed-in view on the right highlights the steric hindrance by AMP-6 of cA_6_ that restricts the movement of R25.

Compared to the structure of apo ApCsm6, cA_5_ binding induces a large structural movement in CARF-1 (Fig. 5d, Supplementary Movie 2). Of note, R25 on the lid helices undergoes an 8.5 Å movement to coordinate the 5’-phosphate of AMP-5 (Fig. 5e and Supplementary Movie 3). The adenosine moiety of AMP-5, which in the cA_6_-bound structure only forms a base-stacking interaction with that of AMP-1, is accommodated within a pocket formed by loop Q561-G570 (Fig. 5d, e). Additionally, AMP-2 binds within a more compact pocket, facilitated by the engagement of loop A59-R70 (Fig. 5d, e), which is disordered in both the apo and cA_6_-bound states. Specifically, F68 on this loop forms π-π stacking with the adenine group of AMP-2 (Fig. 5e). Similarly, AMP-3 engages in an increased number of hydrogen-bonding interactions owing to the binding of loop D493-V501 (Fig. 5d, e).

Structural comparison of the cA_6_- and cA_5_-bound CARF domains reveals a steric clash between the lid helix (containing R25) and AMP-6 during the conformational transition (Fig. 5f), explaining the impaired domain closure upon cA_6_ binding.

Together, these structural insights elucidate the molecular basis for ApCsm6’s preference for cA_5_ over cA_6_ as its activator.

### Mechanism of allosteric activation

Upon cA_5_ binding, CARF-1 undergoes a substantial conformational change toward CARF-2 (Fig. 6a and Supplementary Movie 2). This structural transition propagates to the HEPN domain, inducing conformational changes in the conserved R-X₄₋₆-H motifs, including R295/H369 in HEPN-1 and R723/H730 in HEPN-2 (Fig. 6b). Alanine substitution of any residue in the R-X₄₋₆-H motifs (R295, H369, R723, or H730) completely abolished ApCsm6’s ribonuclease activity (Fig. 6c). In particular, cA_5_ binding triggers a ∼ 90° rotation of R295’s guanidinium group from its apo conformation, where it was stabilized by interactions with D299 (Fig. 6b). This mechanism parallels the activation of TtCsm6 by cA_4_, whose binding releases R415 from E332, positioning its guanidine side chain at the catalytic center ^20^. In addition to R295, residues R723 and H730 also undergo subtle conformational changes following cA_5_ binding (Fig. 6b). These structural adjustments likely optimize the positioning of catalytic residues for RNA substrate engagement, enabling efficient ribonucleolytic cleavage. In contrast, in the cA_6_-bound structure, the conformation of R295 retains its apo-like configurations, and H730 exhibits significant deviation from the catalytic center (Supplentary Fig. 4a), displaying an inactive conformation.

**Fig. 6.**
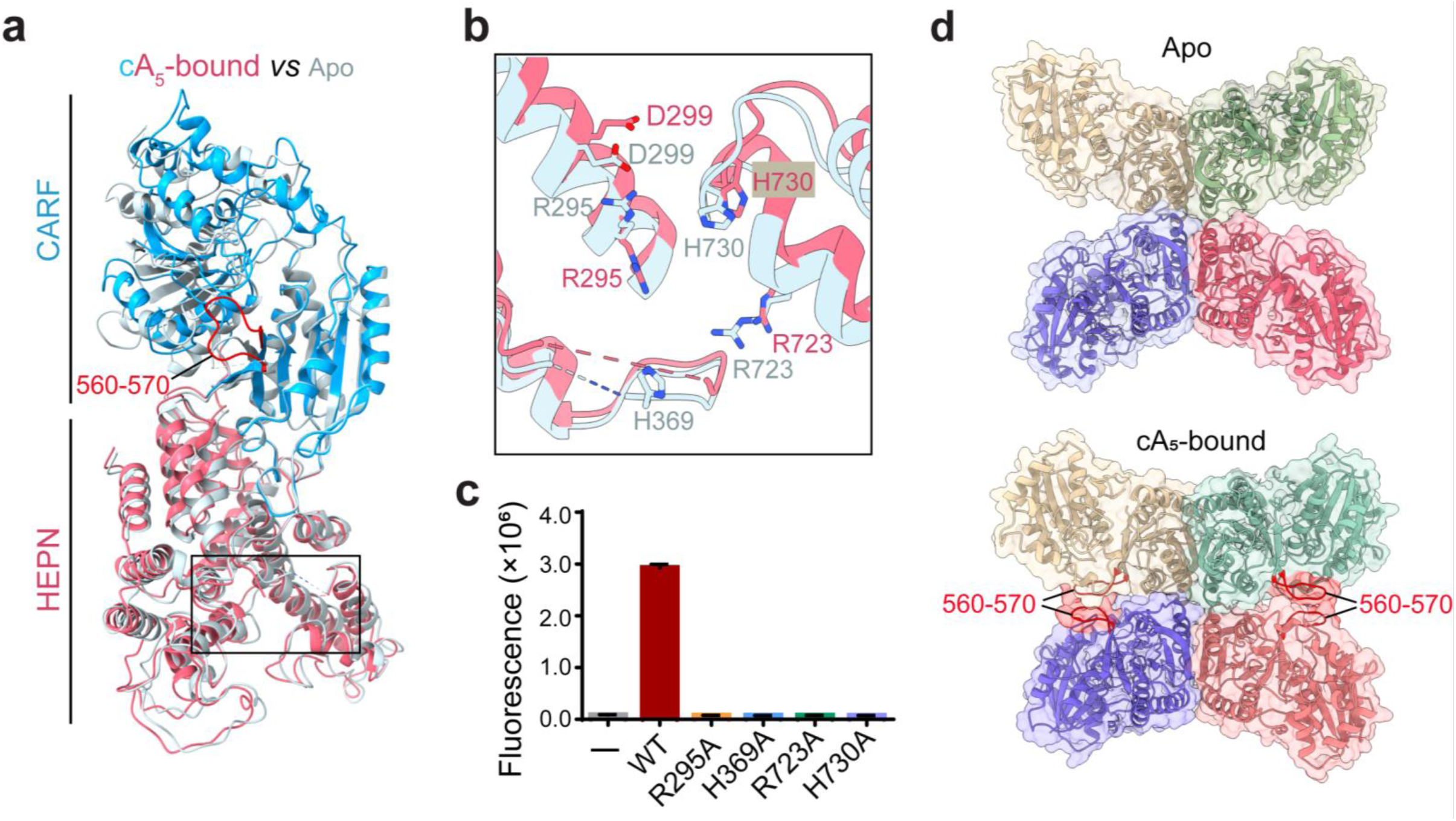
Mechanism of the allosteric activation by cA_5_. a, Structural comparison of apo and cA_5_-bound ApCsm6. The CARF and HEPN domains of cA_5_-bound ApCsm6 are colored blue and red, respectively. Apo ApCsm6 is in light blue. **b,** Close-up view of active site architecture in apo (light blue) and cA_5_-bound (red) ApCsm6. Catalytic residues are shown as sticks. Disordered regions are indicated by dashed lines. **c,** Effect of HEPN domain mutations on the ribonuclease activity of ApCsm6. Values are means ± SD, n = 3 replicates. **d,** Comparison of tetramerization interface of apo and cA_5_-bound ApCsm6. CARF domains from four monomers are shown in distinct colors, overlaid with overlaid with an 80% transparent surface. Loops 560-570 are highlighted in red.

In addition to the changes in HEPN domain, cA_5_ binding also induces additional tetramerization interface of ApCsm6 (Fig. 6d). Structural comparisons reveal that the V560-G570 loop, which remains disordered in both apo and cA_6_-bound states due to inherent flexibility (Fig. 6a and Supplementary Fig. 4a, b), becomes well-ordered upon cA_5_ binding. In this conformation, the loop forms stable interactions with its counterpart in adjacent monomers (Fig. 6a, d), expanding the interfacial area between monomers from 428.8 Å² to 651.1 Å². Since tetramerization is essential for ApCsm6’s ribonuclease activity, these additional intermonomer contacts likely stabilize the catalytically active tetramer.

Taken together, these findings reveal a dual role for cA_5_ in orchestrating both local active site remodeling and global oligomerization dynamics to achieve optimal enzymatic function.

## Dscussion

Type III CRISPR-Cas systems feature the signature Cas10 protein that generates cOA messengers (typically containing 3-6 AMPs) in response to viral mRNA detection ^11^. While numerous cOA-dependent effector proteins have been characterized, it is striking that no cA_5_-dependent effectors have been identified to date, particularly given that cA_5_ is among the most abundant cOA species produced during phage infection ^32^. In this study, we report that Csm6 from the type III CRISPR system of *Actinomyces procaprae* preferentially utilizes cA_5_ as its activator, with over 1,000-fold greater potency compared to cA_6_.

Most characterized Csm6 proteins assemble composite CARF domains through homodimerization and preferentially bind symmetric ligands like cA_4_ and cA_6_ ^10^. Unlike canonical Csm6 proteins, ApCsm6 contains two tandem CARF-HEPN modules (CARF1-HEPN1-CARF2-HEPN2) within a single polypeptide chain, forming an asymmetric binding pocket. This unique architecture accommodates cA_5_ perfectly, with three AMPs bound to CARF-1 and two to CARF-2. Although cA_6_ can bind, only five of its six AMPs are unambiguously positioned in the CARF domain. The sixth AMP exhibits weak electron density and minimal protein interactions. These structural features thus determine ApCsm6’s selective binding preference for cA_5_ in its CARF domain.

Compared to the structure of apo ApCsm6, cA_5_ binding induces a large structural movement of CARF-1 toward CARF-2, converting the CARF domain to a closed conformation (Supplementary Movie 2). The CARF domain conformational changes propagate to the HEPN domain, reorganizing the catalytic R-X₄₋₆-H motif into an active configuration that enables ribonuclease activity. In contrast, cA_6_ binding cannot induce full CARF domain closure due to steric interference between the CARF-1 lid helix and cA_6_’s sixth AMP, explaining its significant weaker activation potency relative to cA_5_. In addition to inducing conformational changes within individual ApCsm6 monomers, cA_5_ but not cA_6_ binding also enhances tetramerization by significantly expanding the oligomerization interface (∼223 Å² increase versus apo state). Since tetramer formation is essential for ApCsm6’s ribonuclease activity, this enhanced interface therefore stabilizes the catalytically active complex.

To prevent excessive RNA cleavage that could induce host cell dormancy or death during early viral infection, many Csm6 proteins act as self-limiting ribonucleases via intrinsic cOA-degrading activity. This activity is typically mediated by their CARF domains ^19,20,22,24,31,32^ or by acquired viral anti-CRISPR ring nucleases ^21,33^. In contrast, the CARF domain of ApCsm6 lacks cOA-cleaving activity, instead, it relies on its HEPN domain to degrade excess cOA. As a consequence, cOA bound within the CARF domain may be protected from degradation, consistent with our observation that pre-incubation of cOA with ApCsm6 did not completely eliminate ApCsm6’s ribonuclease activity (Fig. 1f). The mechanism by which host cells ultimately terminate ApCsm6-cOA signaling is currently unclear. However, it is possible that cells may need to maintain basal-level immunity to provide continuous protection against persistent environmental threats. Future cell-based studies should address this critical regulatory gap in type III CRISPR-mediated immunity.

## Data availability

The cryo-EM density maps have been deposited to the Electron Microscopy Data Bank under accession numbers EMD-65609 (apo ApCsm6), EMD-65610 (ApCsm6-cA_6_) and EMD-65611 (ApCsm6-cA_5_). The corresponding atomic coordinates are available in the RCSB Protein Data Bank under accession codes 9W3U (apo ApCsm6), 9W3V (ApCsm6-cA_6_) and 9W3W (ApCsm6-cA_5_).

## Acknowledgements

Single particle cryo-EM data were collected at the Westlake University Cryo-EM Facility. We thank the Westlake University High-Performance Computing Center for computational resources and technical assistance.

## Author contributions

Ruyi Shi performed protein purification, biochemical analyses and Cryo-EM sample preparation. Mengquan Yang and Yusong Liu performed cryo-EM grid preparation, data collection and processing, under the supervision of Haishan Gao. Haishan Gao performed model building and structural refinement. Zhonghui Lin supervised the project, analyzed the data and wrote the paper with input from all authors.

## Methods

### Protein Expression and Purification

The ApCsm6 cDNA (GenBank ID: WP_136192673) was synthesized by GenScript Corporation (Nanjing, China) after codon optimization and cloned into the pET-28a vector, with an N-terminal His_6_ tag. The pET-28a-ApCsm6 plasmid was transformed into *E. coli* Rosetta (DE3) cells. Protein expression was induced by 0.5 mM isopropyl β-D-1-thiogalactopyranoside (IPTG) overnight at 17°C. Cells were harvested and resuspended in lysis buffer containing 50 mM Tris-HCl (pH 7.5), 300 mM NaCl, 5% glycerol, and 10 mM imidazole. After sonication and centrifugation, the His_6_ - tagged protein was pooled onto Ni-NTA column (Union Biotech, China). After thorough washing, the protein was eluted with lysis buffer supplemented with 200 mM imidazole. The protein was further purified using 15Q and Superdex 200 10/300 GL columns (GE Healthcare Life Sciences), and stored at -80°C in 50 mM Tris-HCl (pH 7.5) and 250 mM NaCl. The expression and purification of ApCsm6 mutants and truncated variants followed the same protocol as described above.

### ssRNA cleavage assay

The ssRNA cleavage activities of ApCsm6 and its variants were assessed using both the denaturing polyacrylamide gel electrophoresis (PAGE) and fluorescence resonance energy transfer (FRET)-based assays as previously described ^20^. For the gel-based assay, 250 nM FAM-labeled ssRNA (5’-ACUGCAACGCAAUAU- ACCAUAGCU-3’) was incubated with 100 nM ApCsm6 and 100 nM cOA (cA_5_ and cA_6_, Biolog Life Science Institute, Germany) at 37°C for 45 min, in cleavage buffer containing 20 mM Tris-HCl (pH7.0), 50 mM KCl, and 25 mM EDTA. Subsequently, the reaction products were separated by a 12% urea denaturing gel, which was imaged using the ChemDoc Touch imaging system (Bio-Rad). For the FRET-based assay, 200 nM synthetic RNA reporter (FAM-UGUUCGACGA-BHQ1) was incubated with 10 nM ApCsm6 and 1 nM cA_5_ (or 200 nM ApCsm6 and 200 nM cA_6_ for cA_6_ activation assay) in cleavage buffer containing 20 mM Tris-HCl (pH 7.0), 50 mM KCl, and 25 mM EDTA. Fluorescence values were monitored by a microplate reader (BioTek) for 2 h at 1-min intervals, with excitation at 490 nm and emission at 520 nm. To investigate ApCsm6 activation by linear oligonucleotides, we adapted the LbuCas13a-Csm6 tandem nuclease assay as previously described ^37^. Briefly, 40 nM LbuCas13a was pre-incubated with 20 nM crRNA in 25 mM HEPES (pH 7.0), 50 mM KCl, 10 mM MgCl₂, and 5% glycerol. A mixture containing 100 nM ApCsm6, 200 nM reporter RNA (FAM-10A-BHQ1), and 2 μM activator RNA (A_2-6_U_6_) was then added. The reaction was initiated by adding 100 pM target RNA complementary to the crRNA spacer region. Fluorescence intensity was monitored using a microplate reader.

### cOA cleavage assay

The cOA cleavage activity of ApCsm6 was assessed by HPLC and MALDI-TOF MS analyses, as previously described ^30^. Briefly, in a 50-μL reaction, 40 μM cOA was incubated with 2 μM ApCsm6 at 37°C for 2 h, in cleavage buffer containing of 20 mM Tris-HCl (pH 7.0) and 50 mM KCl. Reaction products were extracted using an equal volume of chloroform-isopentanol mixture (24:1), and the aqueous phase was collected for further HPLC and MALDI-TOF MS analysis.

HPLC analysis was conducted on a Agilent 1260 Infinity II LC System, equipped with an RX-C18 column (2.1×100 mm, 5 µm) (Zhongpu Science). All components were eluted using a linear gradient of mobile phase A (0.1% trifluoroacetic acid in water) and mobile phase B (0.1% trifluoroacetic acid in acetonitrile) at a column temperature of 40°C. The eluent was monitored by UV detection at 259 nm.

For MALDI-TOF MS analysis, cOA and its cleavage products were mixed with matrix solution, and 1 μL of the mixture was applied onto a 384 MTP AnchorChip. After drying and crystal formation, the chip was transferred to the excitation source for analysis, using a frequency-tripled Nd:YAG (355 nm) laser. The source region (metal probe) was maintained at 20 kV (AUTOFLEX III MALDI-TOF, Bruker Corporation, Germany). Samples were analyzed using FlexControl software, with data processed via FlexAnalysis.

### Cryo-EM data collection and image processing

For cryo-EM grid preparation, 3 μl ApCsm6 samples (∼0.5 mg/ml) were applied onto glow-discharged holey carbon grids (Quantifoil Cu with 2 nm Carbon, R1.2/1.3, 300 mesh), blotted with a Vitrobot Marker IV (Thermo Fisher Scientific) for 3 s under 100% humidity at 4°C, and subjected to plunge freezing into liquid ethane. Cryo-EM data for ApCsm6 (Apo) was collected using the Thermo Fisher Titan Krios microscope at 300 kV equipped with a Gatan K3 Summit direct electron detector (super-resolution mode, at a nominal magnification of 81,000, pixel size 1.087) and a GIF-quantum energy filter; cryo-EM data for ApCsm6-cA_6_ and ApCsm6-cA_5_ were collected using the Thermo Fisher Titan Krios microscope at 300 kV equipped with a Falcon 4i Summit direct electron detector (at a nominal magnification of 130,000, pixel size 0.97) and a GIF-quantum energy filter. Total electron doses were set at 50 e^-^/Å^2^. Defocus values were set from -1.0 to -2.0 μm. EPU (Thermo Fisher) was used for fully automated data collection.

All micrograph stacks were motion corrected with MotionCor2 ^38^ resulting in a pixel size of 1.087 or 0.97 Å, indicated on the flowchart. Contrast transfer function (CTF) parameters were estimated using Gctf. Most steps of image processing were performed using cryoSPARC ^39^.

For 3D processing of apo ApCsm6, 4,304,615 particles were auto-picked from 1,608 micrographs. These particles were extracted with Bin 4 and underwent multiple rounds of reference-free 2D classification. After removing obvious ice contaminants and junk particles, 1,484,826 particles were retained and then re-extracted without binning. Next, ab initio models were constructed and employed for heterogeneous 3D refinement. The resulting class of 149,420 particles was re-extracted, further classified via 2D and ab initio methods, and subjected to Non-Uniform reconstruction for subsequent structural analysis. The overall resolution of the apo ApCsm6 map was determined as 2.59 Å using the Fourier Shell Correlation (FSC) 0.143 criterion ^40^.

For 3D processing of the ApCsm6-cA_6_ dataset, 7,333,325 particles were automatically selected from 5,340 micrographs. Extracted with Bin 4, these particles were subjected to multiple rounds of reference-free 2D classification. Following exclusion of ice contaminants and junk particles, 1,672,108 particles were kept and re-extracted without binning. Ab initio models were then generated and used for heterogeneous 3D refinement. The class containing 667,978 particles was subjected to further non-uniform refinement and local refinement (both with and without D2 symmetry application) for structural analysis. The global resolution of the ApCsm6-cA_6_ map was 2.53 Å based on the FSC 0.143 criterion.

For 3D processing of the ApCsm6-cA_5_ data, 1,969,041 particles were auto-picked from 1,994 micrographs. After extraction with Bin 4, the particles underwent several rounds of reference-free 2D classification. Post-removal of ice contamination and junk particles, 183,599 particles were retained and re-extracted without binning. Ab initio models were built and subsequently utilized for heterogeneous 3D refinement. The class of 143,784 particles was then processed with further non-uniform refinement and local refinement (with and without D2 symmetry) for structural analysis. The overall resolution of the ApCsm6-cA_5_ map was 2.67 Å as per the FSC 0.143 criterion.

### Model building

The structure of ApCsm6 monomer predicted by Alphafold 2 ^41,42^ was used as the starting models and docked into the final EM maps with UCSF Chimera ^43^. The models were manually adjusted and iteratively built in COOT ^44^ and then refined against summed maps using phenix.real_space_refine implemented in PHENIX ^45^ until the validation data were reasonable. FSC values were calculated between the resulting models and the two half-maps, as well as the averaged map of the two half-maps. The quality of the models was evaluated with MolProbity ^46^. The structure validation statistics were listed in table S1. All structural figures were prepared with Chimera X ^47^.

