## Supplementary information for "Mechanistic Insights into Cyclic Penta-Adenylate-Mediated Activation of Type III CRISPR Ribonuclease Csm6"

### Supplementary Figures

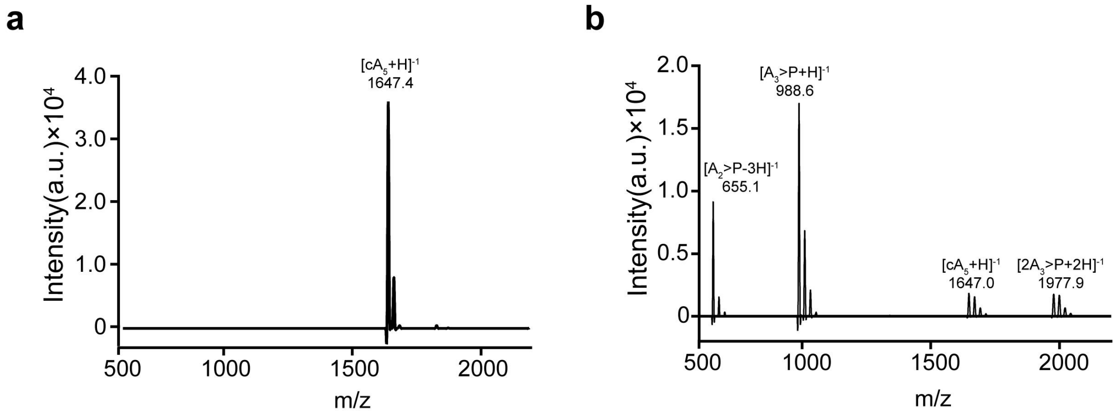


**Supplementary Fig. 1 │ Mass Spectrometry analyses of $cA_5$ cleavage by ApCsm6.** **a,** Mass spectra of $cA_5$. **b,** Mass spectra of the reaction products of $cA_5$. 40 μM $cA_5$ was incubated with 2 μM ApCsm6 at 37°C for 2 h. Oligonucleotides were extracted with chloroform-isopentanol and subjected to MALDI-TOF MS analysis.

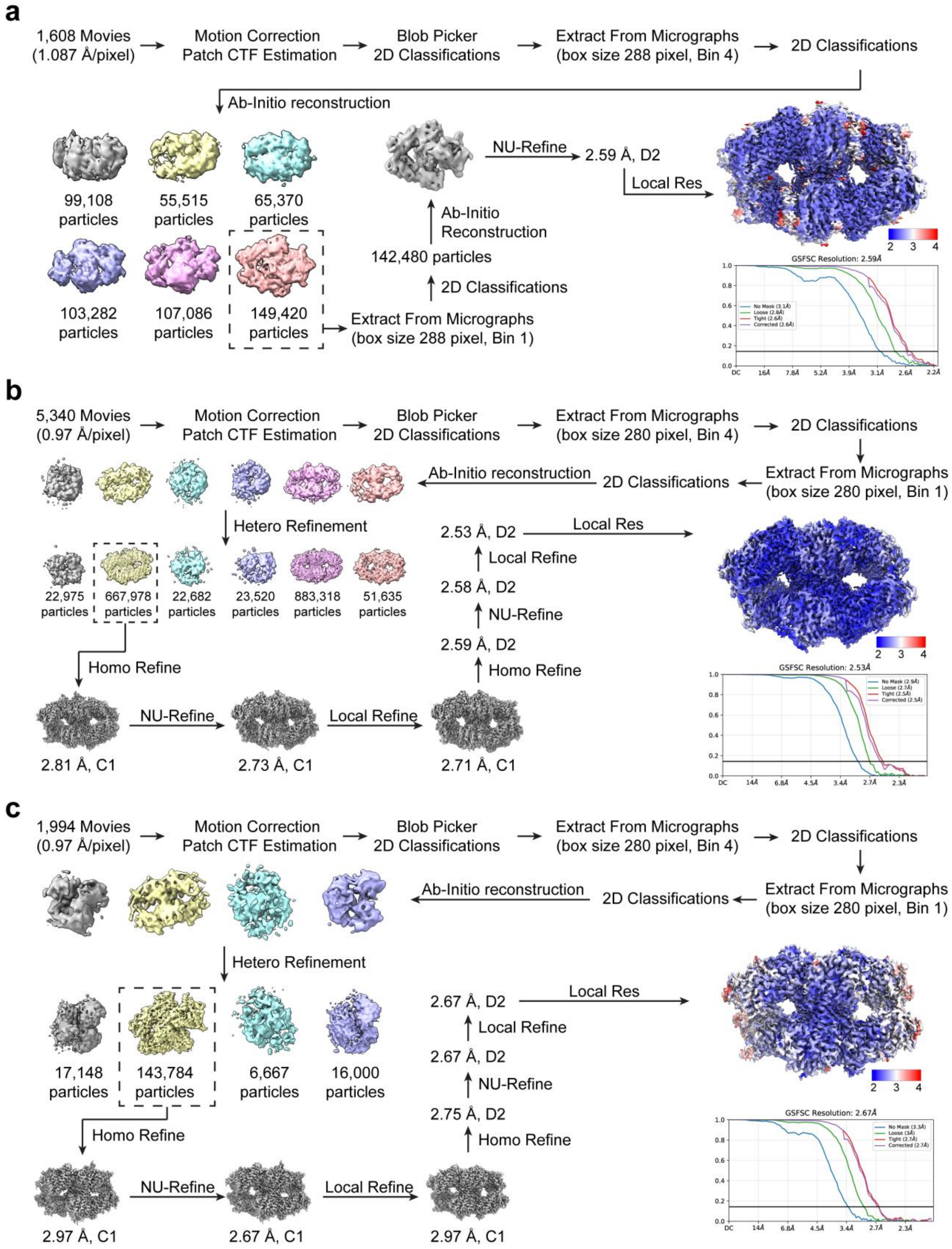


**Supplementary Fig. 2 | Flowchart of cryo-EM data processing. a,** Cryo-EM analysis of ApCsm6. **b,** Cryo-EM analysis of ApCsm6 in complex with $cA_6$. **c,** Cryo-EM analysis of ApCsm6 in complex with $cA_5$.

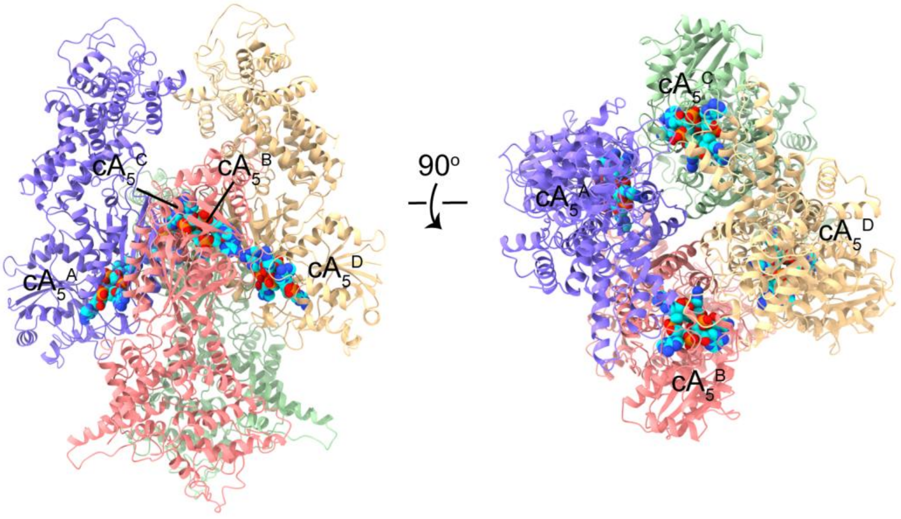


**Supplementary Fig. 3 | Structure of ApCsm6 in complex with cA$_5$.** ApCsm6 is shown in Cartoon representation, and cA$_5$ is depicted as cyan sphere. Each individual ApCsm6 monomer is shown in a distinct color.

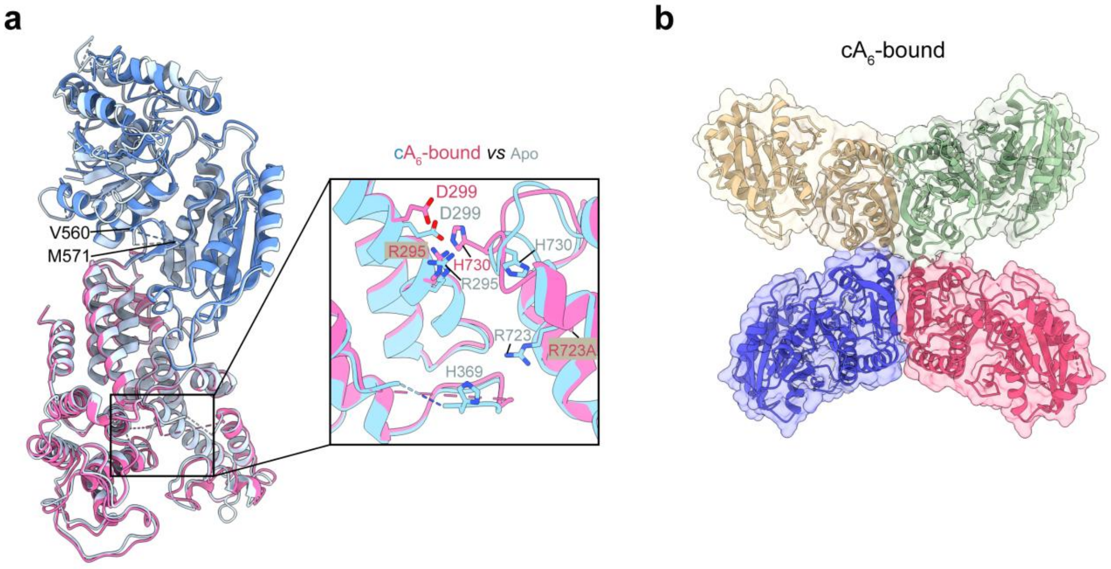


**Supplementary Fig. 4 | Structure of ApCsm6 in complex with cA$_6$. a,** Structural alignment of cA$_6$-bound ApCsm6 with its apo form. The CARF and HEPN domains of cA$_6$-bound ApCsm6 are colored blue and red, respectively. Apo ApCsm6 is in light blue. Right panel: close-up view of active site architecture within the HEPN domain of apo and cA$_6$-bound ApCsm6. Catalytic residues are shown as sticks. Disordered regions are indicated by dashed lines. **b,** Tetramerization interface of cA$_6$-bound ApCsm6. CARF domains from four monomers are shown in distinct colors, overlaid with overlaid with an 80% transparent surface.

**Supplementary Table 1. Cryo-EM data collection, refinement, and validation statistics**

| Complexes | Apo ApCsm6 | ApCsm6-$cA_6$ | ApCsm6-$cA_5$ |
|---|---|---|---|
| PDB ID | 9W3U | 9W3V | 9W3W |
| EMDB ID | EMD-65609 | EMD-65610 | EMD-65611 |
| **Data Collection and Processing** | | | |
| Microscope | FEI Titan Krios | | |
| Voltage | 300 | | |
| Electron dose ($e^-$/Å$^2$) | 50 | | |
| Defocus range (µm) | -1.0 to -2.0 | | |
| Detector | Gatan K3 | Falcon 4i | Falcon 4i |
| Magnification | 81,000 | 130,000 | 130,000 |
| Pixel size (Å/pixel) | 1.087 | 0.97 | 0.97 |
| Micrographs (no.) | 1,608 | 5,340 | 1,994 |
| Final particles (no.) | 149,420 | 667,978 | 143,784 |
| Symmetry imposed | D2 | D2 | D2 |
| Map resolution (Å) | 2.59 | 2.53 | 2.67 |
| FSC threshold | 0.143 | 0.143 | 0.143 |
| **Model composition** | | | |
| Chains | 4 | 8 | 8 |
| Atoms | 22,972 | 22,636 | 24,048 |
| Protein residues | 3,044 | 2,928 | 3,132 |
| Nucleotide | 0 | 24 | 20 |
| Ligands | 0 | 0 | 0 |
| **Refinement** | | | |
| Initial model used | Alphafold 2 | | |
| Model resolution (Å) | 2.8 | 2.8 | 3.0 |
| FSC threshold | 0.143 | 0.143 | 0.143 |
| Map sharpening B factor (Å$^2$) | -106.9 | -101.7 | -94.4 |
| **Validation** | | | |
| MolProbility score | 2.40 | 1.77 | 1.86 |
| Clash score | 7.05 | 3.88 | 4.75 |
| Rotamer outliers (%) | 4.39 | 2.24 | 2.30 |
| $C_\beta$ outliers (%) | 0.36 | 0.00 | 0.04 |
| **R.m.s deviations** | | | |
| Bonds length (Å) | 0.005 | 0.004 | 0.003 |
| Bonds Angle (°) | 0.879 | 0.727 | 0.682 |
| **Ramachandran plot (%)** | | | |
| Favored | 90.73 | 95.24 | 95.11 |
| Allowed | 8.13 | 3.64 | 4.18 |
| Outliers | 1.14 | 1.12 | 0.71 |

**Supplementary Movie 1**: Conformational changes of ApCsm6 upon $cA_6$ binding.

**Supplementary Movie 2**: Conformational changes of ApCsm6 upon $cA_5$ binding.

**Supplementary Movie 3**: Close-up view of changes within the CARF domain upon $cA_5$ binding.
